# Design to Data for mutants of β-glucosidase B from Paenibacillus polymyxa: Y333F, A88E, L219Q, A408H, Y173L, E340S, and Y422F

**DOI:** 10.64898/2026.02.04.703908

**Authors:** Alexandra Maduros, Leila Farinsky, Pagkratios Tagkopoulos, Ashley Vater, Justin B. Siegel

**Affiliations:** Genome Center, University of California, Davis 95616, California, United States; Department of Biochemistry & Molecular Medicine, University of California, Davis 95616, California, United States; Department of Chemistry, University of California, Davis 95616, California, United States

## Abstract

This study explores computational design predictions related to experimental enzyme behavior by analyzing seven single-point mutants of β-glucosidase B (BglB) from *Paenibacillus polymyxa*: Y333F, A88E, L219Q, A408H, Y173L, E340S, and Y422F. Each mutation was modeled using Foldit Standalone, and mutant selections were based on predicted thermodynamic stability changes of interest. Six of the seven mutants in this set yielded soluble, expressed protein. Most variants had similar catalytic efficiency compared to the wild type with one exception. The melting temperatures for most variants were also similar to the wild type. Correlation analysis revealed weak but potentially informative relationships between predicted ΔTSE and (a) thermal stability and (b) catalytic efficiency. These results further support known limitations of TSE score as a tool for single point mutation design and add to a growing dataset being generated to build the next generation of functionally predictive protein models.

## INTRODUCTION

Advances in computational protein modeling have enabled the design of proteins with enhanced stability, novel functionalities, and structures not found in nature.^1-5^ Despite this progress, enzyme design tools remain particularly challenging to develop given these protein’s intrinsically dynamic nature with exquisitely evolved active site chemistries and unpredictable propagating effects of allosteric changes.^6^ The motivation of the larger project, which these data add to, is to improve predictive capabilities by creating a comprehensive dataset compiling kinetic and thermal parameters of proteins (*K*_M_, *k*_cat_, *k*_cat_/*K*_M_, *T*_m_).^7^ The Siegel Lab established the Design to Data (D2D) program, which collects kinetic and thermal stability data for single-point mutations of β-glucosidase B (BglB) in a unified, publicly accessible database.^8^ This nationwide initiative is head-quartered at the University of California, Davis. Nearly a decade ago Carlin *et al*. analyzed the first 100 mutations produced from student contributions and found limited correlations between physics-based model predictions and experimentally-obtained reaction rates. The authors recommended that substantial advances in predictive protein modeling would require larger datasets by some orders of magnitude.^9^

Seven single-point mutations of β-glucosidase B (BglB) from *Paenibacillus polymyxa* (A88E, L219Q, Y333F,Y422F, Y173L, A408H, and E340S) were analyzed in this study. Each mutant was characterized for kinetic activity and thermal stability relative to the wild type. Computationally predicted changes in energy and structurally-guided rational design hypothesis were compared with experimentally determined changes in turnover rate, approximate binding affinity, and melting temperature to identify potential correlations between these parameters.

## METHODS

### Designing and modeling of BglB mutants

The seven BglB mutants were modeled using the Foldit Standalone software, which produces a total system energy (TSE) value from the Rosetta energy score function.^2^ The change in TSE (ΔTSE) relative to the WT was calculated for each mutant, and all mutations selected had ΔTSE scores of ten or less.

### Protein production and purification

Mutants used in this study had been previously produced by the D2D Network with kunkel mutagenesis and verified by sanger sequencing (Genewiz, USA). Sequence-verified BglB mutants cloned into cloned pET29b+ vectors were transformed using previously described methods and then induced with IPTG.^10^ The cells were lysed before being purified using nickel-based metal ion affinity chromatography. The yield of total protein in each purified sample was determined by measuring absorbance at 280 nm using the BioTek® Epoch spectrophotometer. Finally, SDS-PAGE gel was used to analyze the presence and purity of the samples.

### Kinetic efficiency and thermal stability assays

Kinetic activity was assessed using the spectrophotometer plate reader (Epoch) at 420 nm to measure the rate of conversion of the substrate 4-nitrophenyl β-D-glucopyranoside (pNPG) following methods from Carlin et al 2016 with recent modifications described by Li et al 2025.^11,12^ The Michaelis-Menten model was then used to calculate the *K*_M_ and *k*_cat_ of each mutant.^2^

The thermal stability of the mutations was assessed using QuantaStudio™ 3 System, using the Protein Thermal Shift (PTS) kit (Applied BioSystem ®, Thermo Fisher) as described by Huang et al 2020. measuring fluorescence reading from 30°C to 80 °C. The two-state Boltzmann model was used to calculate the *T*_m_, the temperature at which 50% of the protein has unfolded. Pearson correlation coefficient (PCC) analysis was used to explore possible relationships between *T*_m_ and *k*_cat_/*K*_M_ and ΔTSE.^13^

## RESULTS

**Figure 1.**
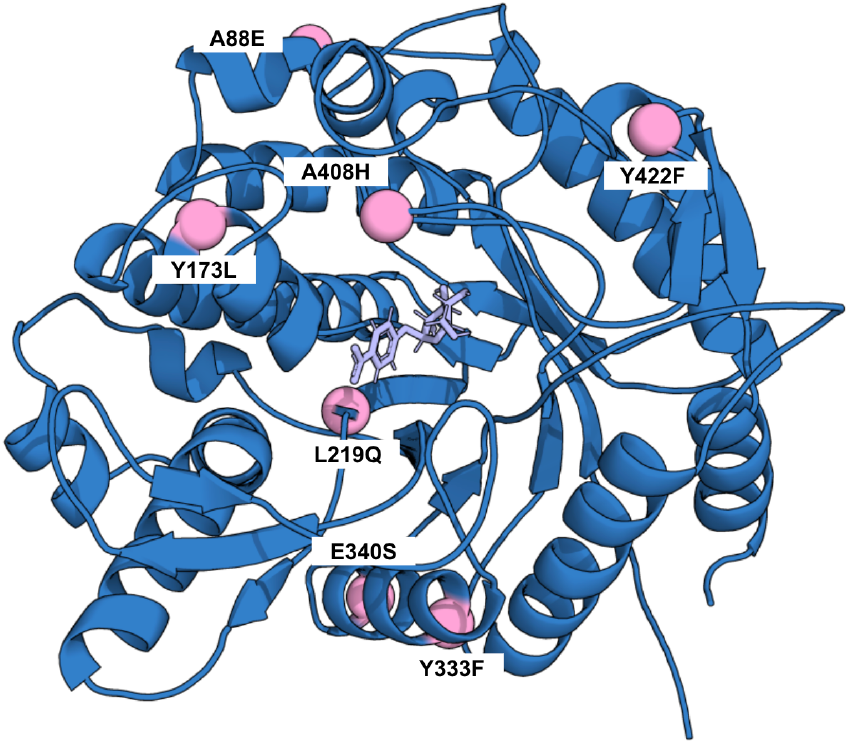
Pymol rendering of BglB enzyme (blue) with substrate 4-nitrophenol beta-D glucopyranoside. The seven mutant locations are highlighted in pink spheres and labeled.

**Figure 2.**
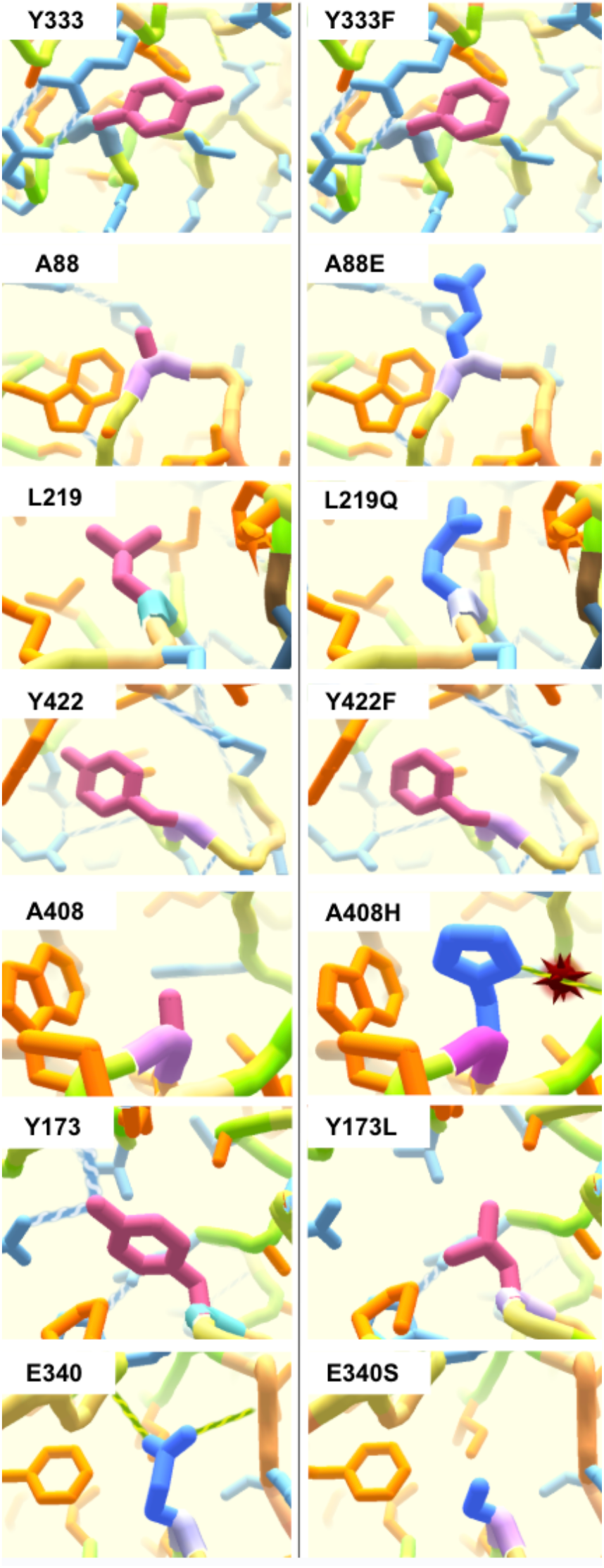
Foldit rendered images of the seven wild type (left) and mutant residues (right), with hydrophobics highlighted in pink (Y333, Y333F, A88, L219, A408, Y173, Y173L, Y422, and Y422F) and hydrophilics/polar highlighted as blue (A88E, L219Q, A408H, E340, and E340S).

### Protein purity and expression

Following IPTG induction protocol, all of the mutants except Y173L expressed and were purified as a soluble protein. The WT and mutants appear in the SDS-PAGE at 50kD bands, indicating the presence and purity of the BglB samples (Figure 3).^11^

**Figure 3.**
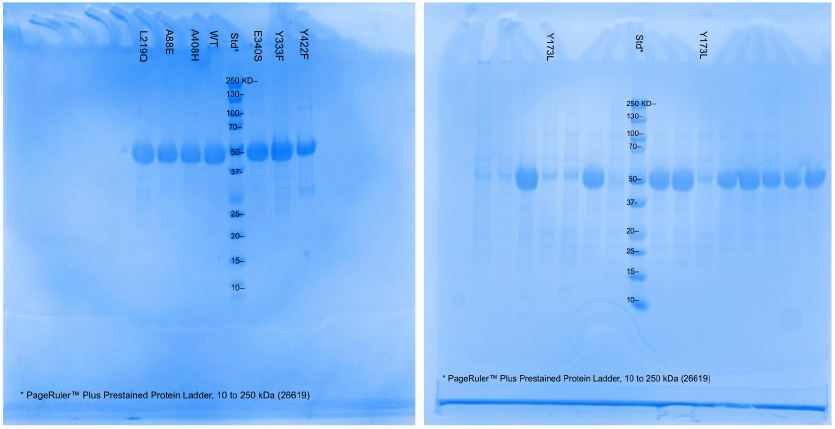
SDS-PAGE analysis of wild type and L219Q, A88E, A408H, E340S, Y333F, Y422F (left) and two samples of Y173L (right). Sample bands formed at approximately 50 kD for WT samples and mutants that expressed, showing BglB purity. Y173L was expressed in biological duplicate, and these samples were run in a second gel (right) to verify the negative expression result; other lanes show samples not described in this study.

### Kinetic activity analysis

The catalytic efficiency was measured by the *k*_cat_ and *K*_M_ values derived by fitting data to the Michaelis-Menten model. The WT’s *k*_cat_/*K*_M_ = 82.96 ± 5.85 mm-1 min−1, which is consistent with previous experimental values in this system.^9,11^ The catalytic efficiency for variant Y422F decreased within ∼4 fold compared to WT (Figure 4), and L219Q showed a small decrease in catalytic efficiency compared to the WT. The catalytic efficiency of A88E increased within ∼2 fold compared to the WT.

**Table 1.** Total system energy (TSE), expression (mg/mL), kinetic parameters (*k*_cat_, *K*_M_, *k*_cat_/*K*_M_), and thermal stability (*T*_m_) of wild-type and seven variant BglB enzymes.

| | TSE | Expression (mg/mL) | $k_{cat}$ (min<sup>-1</sup>) | $K_M$ (mM) | $k_{cat}/K_M$ (mM<sup>-1</sup>min<sup>-1</sup>) | $T_m$ (°C) |
| --- | --- | --- | --- | --- | --- | --- |
| A408H | -1083.6 | 2.60 | $319.5 \pm 6.5$ | $3.4 \pm 0.3$ | $94.3 \pm 8.0$ | $44.8 \pm 0.3$ |
| L219Q | -1083.1 | 3.39 | $305.8 \pm 2.4$ | $5.7 \pm 0.2$ | $53.4 \pm 1.6$ | $45.3 \pm 0.2$ |
| E340S | -1085.3 | 1.80 | $453.0 \pm 9.4$ | $4.2 \pm 0.3$ | $107.4 \pm 8.9$ | $44.5 \pm 0.5$ |
| Y333F | -1089.4 | 2.93 | $355.9 \pm 6.4$ | $4.5 \pm 0.3$ | $78.6 \pm 5.6$ | $43.6 \pm 0.4$ |
| A88E | -1091.1 | 1.18 | $449.3 \pm 19.5$ | $2.3 \pm 0.4$ | $196.3 \pm 37.6$ | $44.4 \pm 0.2$ |
| Y422F | -1090.0 | 2.84 | $146.0 \pm 5.6$ | $6.6 \pm 0.9$ | $22.3 \pm 3.2$ | $46.8 \pm 0.2$ |
| Y173L | -1079.6 | 0.17 | N/A | N/A | N/A | N/A |
| Y173L | -1079.6 | 0.03 | N/A | N/A | N/A | N/A |
| WT | -1089.7 | 3.18 | $375.1 \pm 6.7$ | $4.5 \pm 0.3$ | $83.0 \pm 5.9$ | $44.8 \pm 0.2$ |

**Figure 4.**
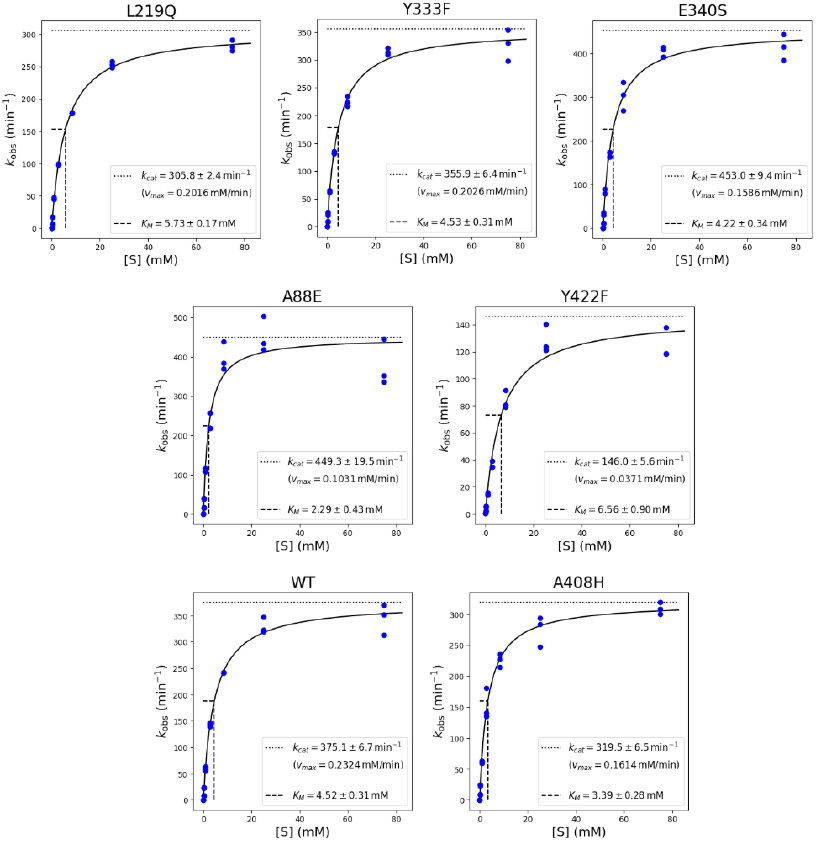
Michaelis-Menten graphs of the WT BglB sample and six tested mutants. Concentration of pNPG was plotted on the horizontal axis and k_obs_ on the vertical axis. The line of best fit for triplicate data is shown in each graph.

### Thermal stability assay

The thermal stability was measured by collecting the triplicated thermal shift assay data from each mutant. The *T*_m_ for the WT was 44.8 ± 0.2°C, similar to previous experimental values in this system.^2^ The thermal stability values for A408H (44.8 ± 0.3°C), A88E (44.4 ± 0.2°C), and E340S (44.5 ± 0.5°C) were within error of WT average, showing no change in thermal stability. Mutant Y422F (46.8 ± 0.2°C) *T*_m_ was about 2°C greater than the WT, and mutant Y333F’s *T*_m_ (43.6 ± 0.4°C) decreased about 1°C compared to the WT.

### Relationships between Foldit TSE Score and functional properties

The relationship between ΔTSE and each of the experimental parameters was analyzed using the Pearson correlation coefficient (PCC). The ΔTSE and catalytic activity (Δ*k*_cat_/*K*_M_) had a PCC of 0.29, while ΔTSE and Δ*T*_m_ had a PCC of 0.01 as shown in Figures 5 and 6, showing weak and no evidence of linear relationship between the investigated parameters, respectively.

**Figure 5.**
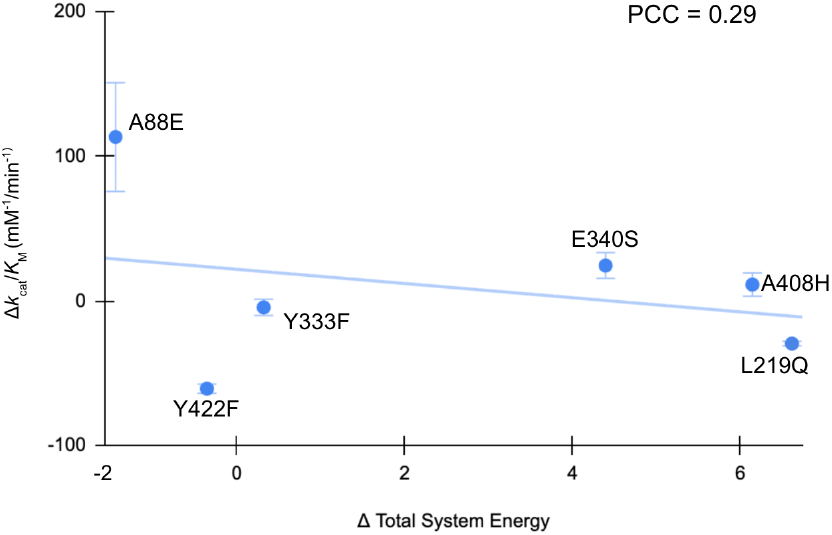
Distribution of Δ*k*_cat_/*K*_M_ values in mM^-1^min^-1^ for each tested mutant plotted against the ΔTSE score predicted by Foldit. The horizontal axis shows ΔTSE and the vertical axis shows mM^-1^min^-1^.

**Figure 6.**
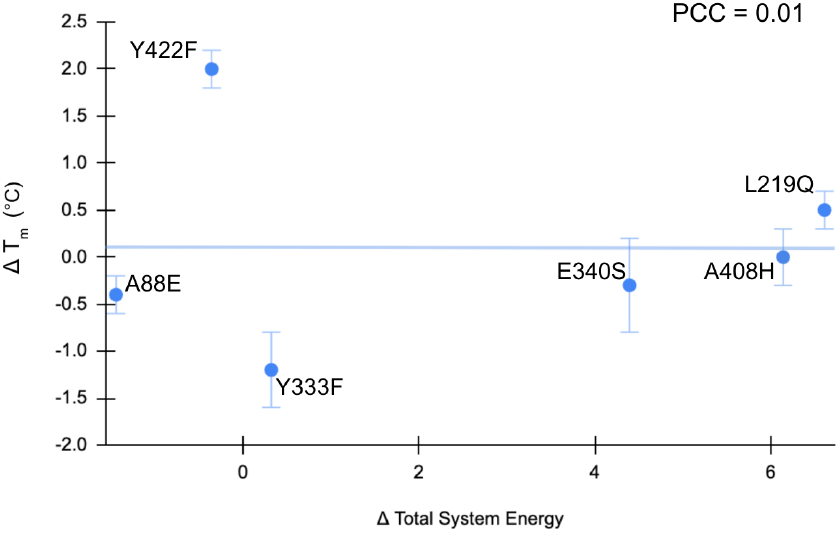
Distribution of *T*_m_ values in degrees Celsius for each tested mutant plotted against the ΔTSE score predicted by Foldit. The horizontal axis shows ΔTSE and the vertical axis shows degrees Celsius.

## DISCUSSION

Across this small set of BglB variants, we observed mixed evidence that local structural features—particularly hydrogen-bond changes and mutation location—track with expression and catalytic performance. These patterns are suggestive but not conclusive given the limited sample size.

### Hydrogen bonding and expression

One notable result was the apparent association between loss of hydrogen bonding and impaired expression. Y173L, the only variant that failed to express, was predicted to lose two hydrogen bonds; this non-expression was reproduced across two independent expression attempts. However, this relationship was not consistent across the dataset: E340S was also predicted to lose two hydrogen bonds yet expressed and performed similarly to WT. Together, these observations suggest that hydrogen-bond loss alone is insufficient to predict expression outcomes and likely depends on the broader structural context of the mutation.

### Proximity to the active site and kinetic effects

We also hypothesized that proximity to the active site would be associated with larger functional perturbations. A408H and L219Q were the closest substitutions to the active site, whereas the remaining mutations were located more peripherally. Most peripheral variants (Y333F, E340S, and A88E) showed no meaningful change in catalytic efficiency, while Y173L failed to express. Among the remaining variants, Y422F was the only mutant with a clear reduction in kinetic efficiency. Overall, active-site proximity did not uniformly predict catalytic disruption in this dataset.

### Relationship between stability metrics and ΔTSE

This small dataset showed essentially no correlation between ΔT_m_ and ΔTSE and a weak negative correlation between Δ*k*_cat_/*K*_M_ and ΔTSE. Given the limitations of the size of this dataset, these analyses should be interpreted cautiously. Notably, other larger studies on D2D data have shown some signal in the association between ΔT_m_ and ΔTSE than between Δ*k*_cat_/*K*_M_ and ΔTSE.^13^

## Conclusions and next steps

In summary, this study expands the BglB D2D dataset by adding expression outcomes and kinetic and thermal measurements for a set of seven single-point mutants, integrated with structural/computational features (e.g., Foldit-derived energetic metrics and predicted hydrogen-bond changes). While these data as an isolated set are underpowered for definitive inference, the results provide initial, testable signals about which features may be informative—particularly the context dependence of hydrogen-bond disruptions for expression and the limited predictive value of mutation proximity to the active site in this variant set. Data that captures many more systematically characterized, targeted sites may improve the accuracy and reliability of predictive models for enzyme design. We are optimistic that continued generation of high-quality, functionally relevant measurements in the D2D dataset—following the ones presented in this study— will strengthen these data-driven approaches and thereby accelerate new enzyme-enabled technologies.

## Acknowledgments

This work was supported by the University of California Davis, the National Science Foundation (award nos. # 2315767, 2118138 and 1827246). The content is solely the responsibility of the authors and does not necessarily represent the official views of the National Science Foundation, or UC Davis. Further, this work references and is contextualized by the Design to Data (D2D) dataset, which has been built through contributions of over 1000 undergraduate students over the past decade.

## Notes

### Competing Interest Statement

The authors have declared no competing interest.

